# Interplay of size and shape in miniaturized land snails

**DOI:** 10.1101/275636

**Authors:** Aydin Örstan

## Abstract

The distribution of the mean greater shell dimensions of 2446 genera of stylommatophoran land snails consists of two groups peaking at 3 mm and 15.3 mm. The 3-mm group includes the miniature snails whose adult shell dimensions range from almost 0.8 mm to less than 1.5 mm. Relative surface area, shell thickness and egg size are discussed as potential factors that may limit the minimum shell dimensions. To obtain uniform distributions, the shell shape is expressed as the expanse angle, defined as the apex angle of a right triangle the height and the base of which are the shell height and diameter, respectively. In terms of expanse angles there are three shell shapes: tall (< 40°), flat (> 50°) and roughly equiaxial (40°-50°). The variation of shell shape with size was analyzed within the morphospace they form. At shell dimensions above 10 mm the lowest expanse angle is about 10°. Below 10 mm, the lower limit of the expanse angle increases as shells get smaller. As a result, no miniature species has a tall shell. It is proposed that two evolutionary functional constraints render small and narrow shells of miniaturized snails nonadaptive. These are the requirements to reduce surface areas to decrease water loss and to have enough volume and a wide enough body whorl to accommodate at least one egg.

## INTRODUCTION

This is a study of the dimensions and shapes of the shells of the smallest (miniaturized) stylommatophoran land snails (Pulmonata: Stylommatophora). I am defining miniaturization as the evolutionarily reduction of body size in some members of an animal taxon to a value at least an order of magnitude below the median body size of the entire taxon. Miniaturization and the accompanying morphological changes have been studied in several animal groups (Hanken & Wake, 1993; Polilov, 2015). Somewhat surprisingly, the miniaturization of land snails has received almost no attention. This is despite the fact that the smallest stylommatophoran land snails, with shell dimensions below 1 mm (Pall-Gergely et al., 2015), are more than two orders of magnitude smaller than the largest species exceeding 200 mm (Winter, 1997) and more than an order of magnitude smaller than the median shell dimension (13.5 mm) of all genera (Fig. 1).

**Figure 1.**
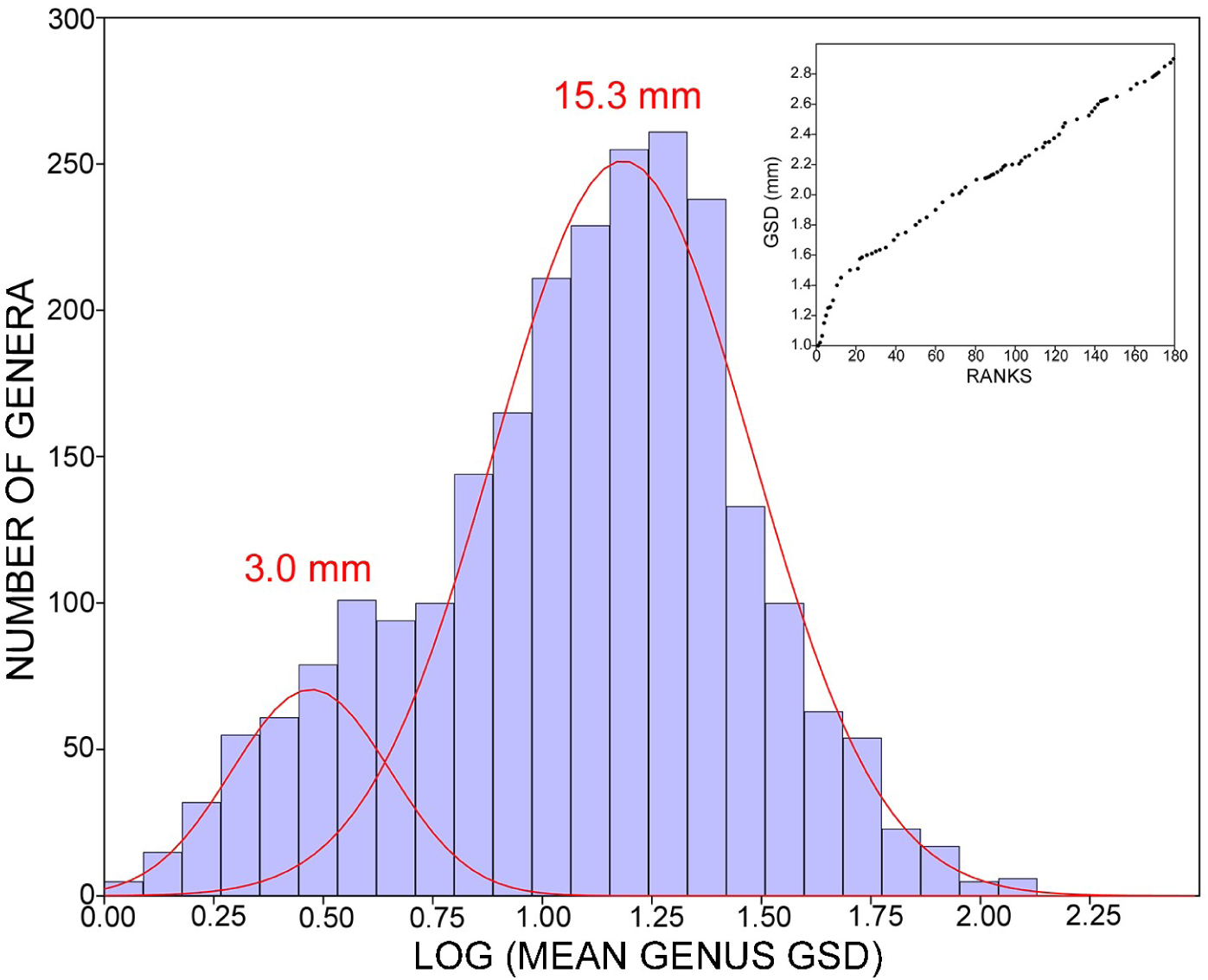
Histogram of the log-transformed mean GSDs of 2446 stylommatophoran genera. The two underlying distributions (red) were resolved using mixture analysis; the back-transformed values of their means were 3.0 mm and 15.3 mm. Inset: Rank plot of the mean GSDs for the first 180 ranks. The abrupt change in slope is at 1.5 mm.

Snail shells have long been characterized by two primary measurements, height (length) and diameter (width). Although it is obviously simplistic to reduce the complicated shape of a shell to two measurements, the ratio of height to diameter (spire index) is a useful proxy for the overall shell shape, especially when comparing the adult shells of closely related species. In his seminal paper, Cain (1977) showed that for stylommatophoran land snails plots of height to diameter displayed a bimodal distribution corresponding to tall and flat shells with a notable paucity of approximately equiaxial shells.

Cain’s (1977) interest was mainly in the comparison of the spire index distributions of the faunas of different geographical areas. To my knowledge, a study of the shell dimensions and shapes of all suitable stylommatophoran genera has not been published. To initiate such a study, I have compiled the shell dimensions of more than 2,400 stylommatophoran genera and subgenera. I then extracted from this compilation mean shell shapes and sizes and analyzed their distribution within the morphospace they form. I present and discuss here only those results of my analysis that apply to the miniaturization of stylommatophoran land snails.

## MATERIALS AND METHODS

I have created a database of mean shell heights (H) and diameters (D) for all of the extant genera and subgenera (hereafter, all taxa are referred to as genera) of stylommatophoran land snails. Slugs and semi-slugs with very small or internalized shells were excluded. This is in line with Cain’s (1977) study, which included only those species that can fully withdraw into their shells. The starting source for my compilation was the 15-part *Treatise* of Schileyko (1998-2007). As my sole purpose was to compile shell dimensions within an already existing taxonomic system, I have used the taxonomy and the dimensions in Schileyko with only a few changes. The current database has 2446 genera. Some of the taxonomic assignments and the dimensions may be revised in the future. However, because the database consists of such a large number of entries, refinements of the dimensions of even a substantial fraction of the genera are unlikely to result in significant modifications in the generalizations presented here.

If one were working with only a few genera each comprising a few species, the genus means could be calculated from the species means. However, when 2446 genera, some of which contain tens of species, are being studied, compilations of dimensions for the individual species and the subsequent calculations would be very time consuming. Therefore, I opted for the most practical method and calculated the mean height and the mean diameter for each genus by averaging the minimum and the maximum values of height and diameter Schileyko (1998-2007) gave for each genus.

Arguably, shell volume, rather than height or diameter, would be a better measure of the overall shell size. However, although the volumes of snail shells larger than about 10 mm in height or diameter can be measured quite accurately (Örstan, 2011), there is no practical method to measure the volumes of smaller shells. And even if one could such measure volumes directly, doing so for a representative shell for each of the 2446 genera would be an arduous undertaking. Calculations using the volume formulae for geometric solids give approximations. Therefore, as a practical proxy for the overall shell size, I have used the greater shell dimension (GSD), which is the greater of the mean height or the mean diameter for a genus (or the greater of the height or the diameter for a single shell). The mixture analysis of the distribution of the GSDs of all genera (Fig. 1) was done with PAST 3.16 (https://folk.uio.no/ohammer/past/). Because the analysis method expects the data to be from a mixture of normally distributed populations, the GSDs were transformed logarithmically to correct for the right skew in their distribution. The shells of *Punctum smithi* Morrison and *Guppya sterki* (Dall) (Table 2), both from Frederick County, Maryland, U.S.A., were measured with a stereomicroscope using a calibrated reticle. The dimensions of the type of *Carychium nannodes* Clapp (Fig. 3) were from Pilsbry (1948). The dimensions of the shells of *Angustopila* species (Fig. 3) were from Páll-Gergely et al., (2015) and Páll-Gergely (personal communication).

Early in the study, I noticed that the plot of H against D, as in Cain (1977), obscured fine patterns in the distribution of the data. Also, the spire index, being a ratio, had an uneven linear distribution when plotted in the form of a histogram or in a scatter plot against GSD. To overcome these problems, I have visualized the spire index (H/D) as the cotangent of the apex angle of a right triangle the height of which is equal to the shell height and the base is to the shell diameter. Upon taking the inverse cotangent of H/D (or the inverse tangent of D/H, a function more readily available in most calculators) one obtains the apex angle itself. I have dubbed this angle the *expanse angle* (ε) as it is a measure of the spread of a shell along one dimension (Table 1). Geometrically, there are three shell shape groups: tall (< 40°), flat (> 50°) and roughly equiaxial (40°-50°). The expanse angles in degrees form a uniform linear distribution. Therefore, I will present my results and discussion in terms of expanse angles.

**Table 1.** Relationships of shell height (H), diameter (D), greater shell dimension (GSD) and expanse angle (ε) to each other.

| H, D | GSD | $\epsilon$ |
| --- | --- | --- |
| H = D | H or D | 45° |
| H > D | H | < 45° |
| D > H | D | > 45° |

## RESULTS AND DISCUSSION

### Distribution of the shell sizes

The histogram of the log-transformed mean GSDs of 2446 stylommatophoran genera appears to be bimodal with one primary group encompassing most taxa and a smaller group partially buried in the left-hand tail of the former (Fig. 1). The median of the entire distribution is 13.5 mm. I used mixture analysis to try to resolve the histogram into additional groups that may be hidden especially within the primary group. However, the differences in the likelihood values obtained for the models with two, three or four groups were too small to allow for the confident selection of one model over the others. For the two-group model, the means of the resolved groups were 3.0 mm and 15.3 mm (Fig. 1). What is important for the present discussion is that in the three-and four-group models there was also a group with a mean located between 2 mm to 3.6 mm. It appears that, regardless of how many other size groups there may actually be, the smaller genera form a distinct group with a mean GSD of about 3 mm. These genera, representing more than 12 families, form a phylogenetically heterogeneous group. This indicates that small shells have evolved independently and converged towards a GSD of 3 mm.

### Miniature snails

The plot of mean genus GSDs against their ranks by size rises with a steep slope up to the GSD of 1.5 mm. Above that value, the slope decreases sharply to a roughly constant value and the curve continues almost linearly (Fig. 1, inset). I have taken this break point in the continuity of the slope, arbitrary though it may be, as a convenient upper limit to designate the smallest taxa. There are 13 genera with a mean GSD below 1.5 mm. They represent the left-hand tail of the group with the mean genus GSD of 3 mm (Fig. 1). These 13 genera also form a phylogenetically varied group with representatives from eight families.

I now move from genera to individual species whose largest measured adults are less than 1.5 mm in GSD. These are the “miniature” snails. Table 2 lists some miniature snails, including the smallest known species. This is a partial list; to avoid a phylogenetic bias, I included only two *Angustopila* species and excluded the other species smaller than 1.5 mm in *Angustopila* (Pall-Gergely et al., 2015), *Afrodonta* (Solem, 1970) and *Punctum* (Pilsbry, 1948). The purpose of Table 2 is to set approximate limits for shell dimensions and to demonstrate the general trend in the values of the expanse angles of miniature snails. The GSD of the smallest measured adult specimen of the smallest species, *A. subelevata*, is 0.82 mm (Pall-Gergely et al., 2017). This value is probably very close to the lower limit for the adult GSD of stylommatophoran land snails.

**Table 2.**
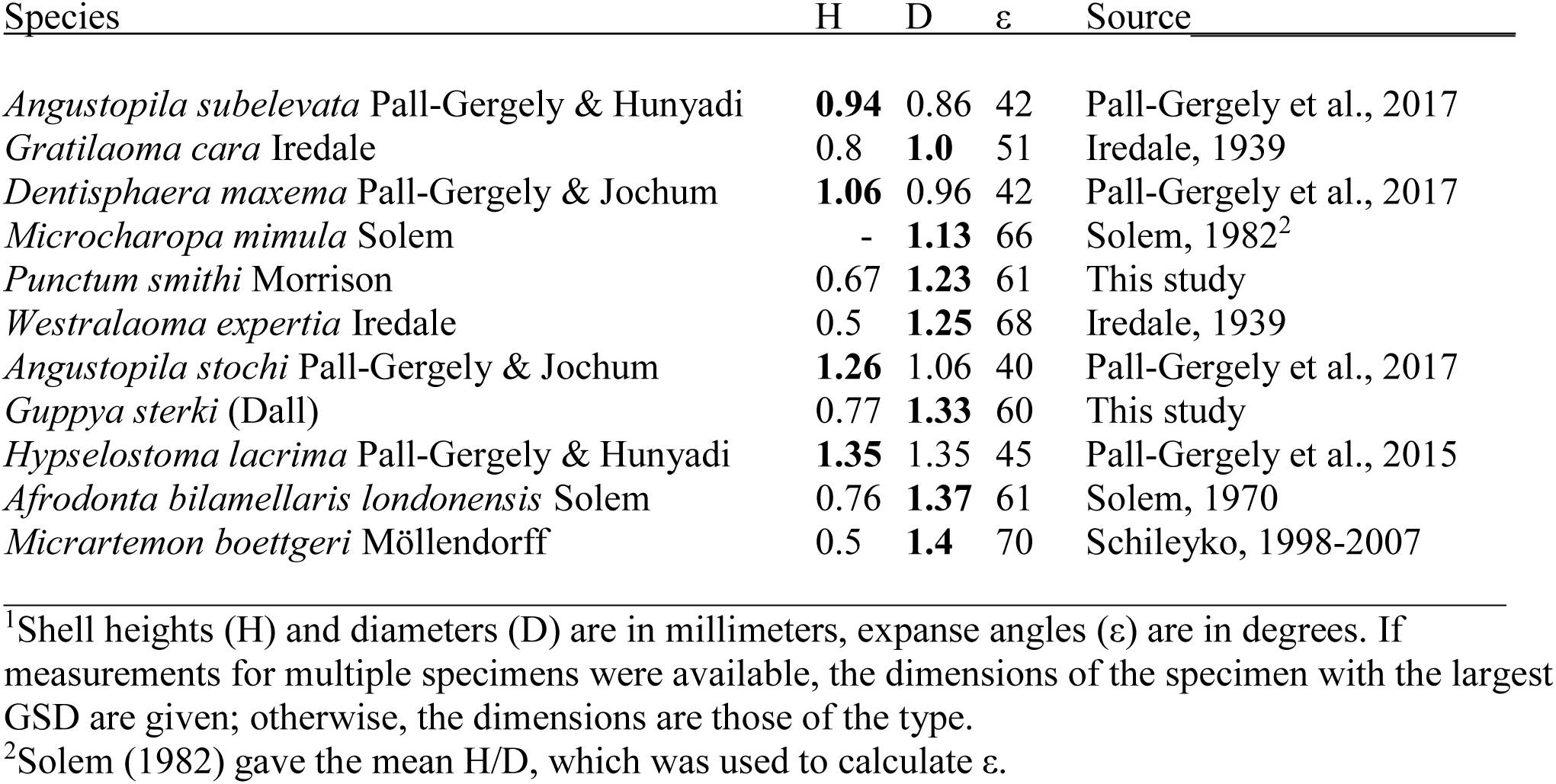
Some of the miniature stylommatophoran land snail species with GSDs (bold face) less than 1.5 mm^1^.

| Species | H | D | $\varepsilon$ | Source |
| --- | --- | --- | --- | --- |
| *Angustopila subelevata* Pall-Gergely & Hunyadi | **0.94** | 0.86 | 42 | Pall-Gergely et al., 2017 |
| *Gratilaoma cara* Iredale | 0.8 | **1.0** | 51 | Iredale, 1939 |
| *Dentisphaera maxema* Pall-Gergely & Jochum | **1.06** | 0.96 | 42 | Pall-Gergely et al., 2017 |
| *Microcharopa mimula* Solem | - | **1.13** | 66 | Solem, 1982<sup>2</sup> |
| *Punctum smithi* Morrison | 0.67 | **1.23** | 61 | This study |
| *Westralaoma expertia* Iredale | 0.5 | **1.25** | 68 | Iredale, 1939 |
| *Angustopila stochi* Pall-Gergely & Jochum | **1.26** | 1.06 | 40 | Pall-Gergely et al., 2017 |
| *Guppya sterki* (Dall) | 0.77 | **1.33** | 60 | This study |
| *Hypselostoma lacrima* Pall-Gergely & Hunyadi | **1.35** | 1.35 | 45 | Pall-Gergely et al., 2015 |
| *Afrodonta bilamellaris londonensis* Solem | 0.76 | **1.37** | 61 | Solem, 1970 |
| *Micrartemon boettgeri* Möllendorff | 0.5 | **1.4** | 70 | Schileyko, 1998-2007 |

<sup>1</sup>Shell heights (H) and diameters (D) are in millimeters, expanse angles ( $\varepsilon$ ) are in degrees. If measurements for multiple specimens were available, the dimensions of the specimen with the largest GSD are given; otherwise, the dimensions are those of the type.
<sup>2</sup>Solem (1982) gave the mean H/D, which was used to calculate $\varepsilon$ .

It should be kept in mind that there are unavoidable errors associated with the dimensions in Table 2. Different observers using different methods may obtain slightly different measurements for a given shell. Moreover, a claim that a certain shell is the smallest (or the largest) of a species would be difficult to justify statistically; the possible existence of an even smaller (or a larger shell) in the wild could never be ruled out. Also, size limits cannot be set easily for the shells of species with indeterminate growth (for example, *Punctum* and *Guppya*).

### Factors influencing the dimensions of miniature snails

A snail’s shell is an integral component of its body. The body and the shell cannot be separated and their dimensions, at least for those snails that can withdraw fully into their shells, cannot evolve independently of each other. As a snail becomes smaller, geometric relations dictate that the surface areas of its body and shell relative to their volumes increase (Schmidt-Nielsen, 1984). Large relative surface areas may have some advantages, including better adhesion and locomotion with the sole of the foot and more efficient gas exchange in the lung. Additionally, a larger shell surface area could accumulate more dew (per volume) under certain atmospheric conditions, which may help snails conserve water (Örstan, 2010). On the other hand, a significant disadvantageous outcome of having large relative surface areas would be increased water loss from the exposed body surfaces and also through the shell, especially if the shell became very thin (Machin, 1967). Relative surface area also depends on shape. Thus, as I will discuss later, the changes in shell shapes observed during miniaturization may, in part, be to reduce evaporation by decreasing relative surface areas.

Shell thickness is probably one factor that limits the minimum size of snail shells. A shell can’t be small and have a thick shell at the same time; otherwise, the volume inside the shell would become insufficient for the snail’s body and the other components (Appendix). Therefore, as a shell gets smaller, its absolute thickness must decrease. The shell may eventually become too thin and fragile to support and protect the snail’s viscera within it against external physical and biological dangers. Decreased shell thickness could also result in increased water loss from the snail’s body (Machin, 1967). Miniature snails may reduce their water loss by confining their activities to high-humidity habitats. However, being in such habitats may increase snails’ risk of getting entrapped in water films and drops if they are not strong enough to overcome the surface tension (see, for example, Morton, 1954). Egg size, an additional limiting factor of snail dimensions, will be discussed later.

### Distribution of the shell shapes

A theoretical morphospace is a two or more dimensional representation of all or a subset of the morphologies potentially available to an organism. Being a theoretical construct, such a morphospace need not include information on the actual prevalence of any of the morphologies in nature. However, once the theoretical morphologies are delimited within the boundaries set by the existing morphologies, one can then search for the evolutionary constraints that may have been responsible for those boundaries as well as the abundances of the existing morphologies (McGhee, 2007).

Absolute size is usually not included in morphospaces (McGhee, 2007). Because I am interested not only in shape but also size, I have constructed a modified morphospace by plotting a measure of shell shape (ε) against size (GSD) (Fig. 2). The horizontal limit of the morphospace is set by the highest mean genus GSD (135 mm). Within this morphospace the distribution of the expanse angles of 2446 stylommatophoran genera separate into two major zones. The zone of the tall shells extends from 10° to 35°, while that of the flat shells from 50° to 75°. The intermediate zone of shell shapes extending from 35° to 50° is less densely populated. This much is in general agreement with Cain’s (1977) findings. However, a closer look hints at the presence of additional clusters, especially within the zone of the tall shells. Also, contrary to the impression one gets from Cain’s (1977) plots of H against D that exactly equiaxial (ε = 45°) shells are rare, the rarest shell shape is actually at about 40° (Fig. 2). A more detailed analysis of the distribution of the expanse angles over all GSD values will be the subject of another study. Here, I will briefly discuss the upper and the lower boundaries of the expanse angle before concentrating on the shell shapes of miniature snails.

**Figure 2.**
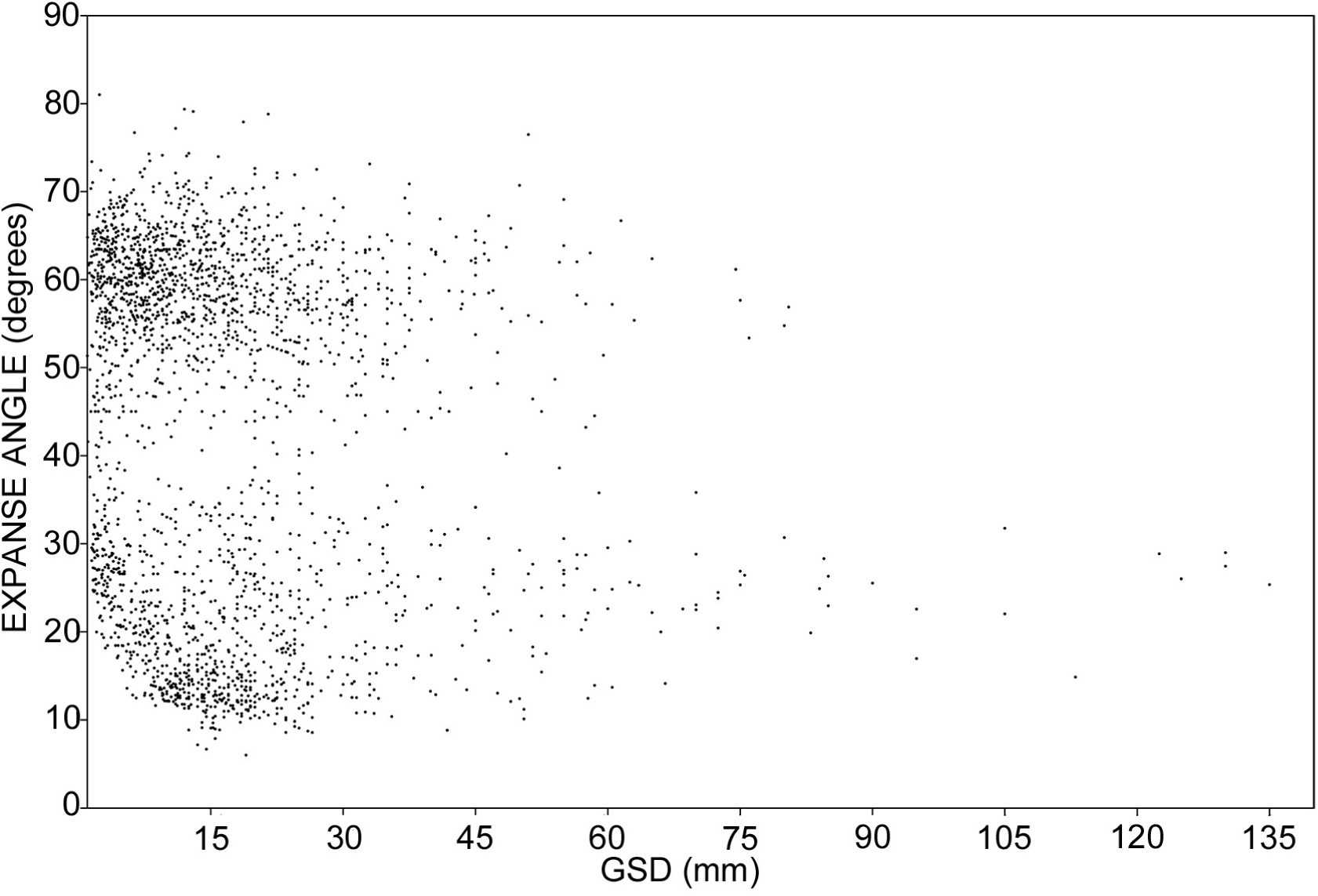
Morphospace of shell shape versus size, expressed as the expanse angle and the mean GSD, respectively, of 2446 stylommatophoran genera.

The upper boundary of the expanse angle is probably at about 75°. Only eight genera have expanse angles larger than 75° and their scatter suggests that they may be outliers arising from limited or erroneous data (Fig. 2). The lower boundary is set by the lowest permissible angle (LPA). Twenty-one genera have expanse angles less than 10°. Fifteen of those genera are in the Urocoptidae. The urocoptid subgenus *Gongylostomella Gongylostomella* (GSD, 19 mm) has the lowest expanse angle of 6° and stands out as a possible outlier (Fig. 2). Many urocoptids have unusual shell shapes with decollated apexes and detached, elongated or laterally projecting body whorls. It is unclear how the dimensions, especially the diameters, of such shells should be measured to obtain numbers comparable to those of more conventional shell shapes. Moreover, many urocoptid dimensions in Schileyko (1998-2007) and elsewhere are of the type specimens only. Future dimensional data from larger samples may eliminate the outliers in Fig. 2. For the time being, if the expanse angles of these 21 genera are ignored, the LPA may be taken to be 10° at GSD values above 10 mm (Fig. 2).

As far as the smaller genera are concerned, the most noteworthy pattern in the distribution of the expanse angles is that below the GSD of 10 mm, the LPA begins to increase along a more or less uniform curve (Figs. 2 and 3). The expanse angles of the individual species in Table 2 do not deviate from the genus means (Fig. 3). The miniature species have either roughly equiaxial (40 ≤ ε ≤50°) or flat (ε > 50°) shells. None has a tall (ε < 40°) shell.

**Figure 3.**
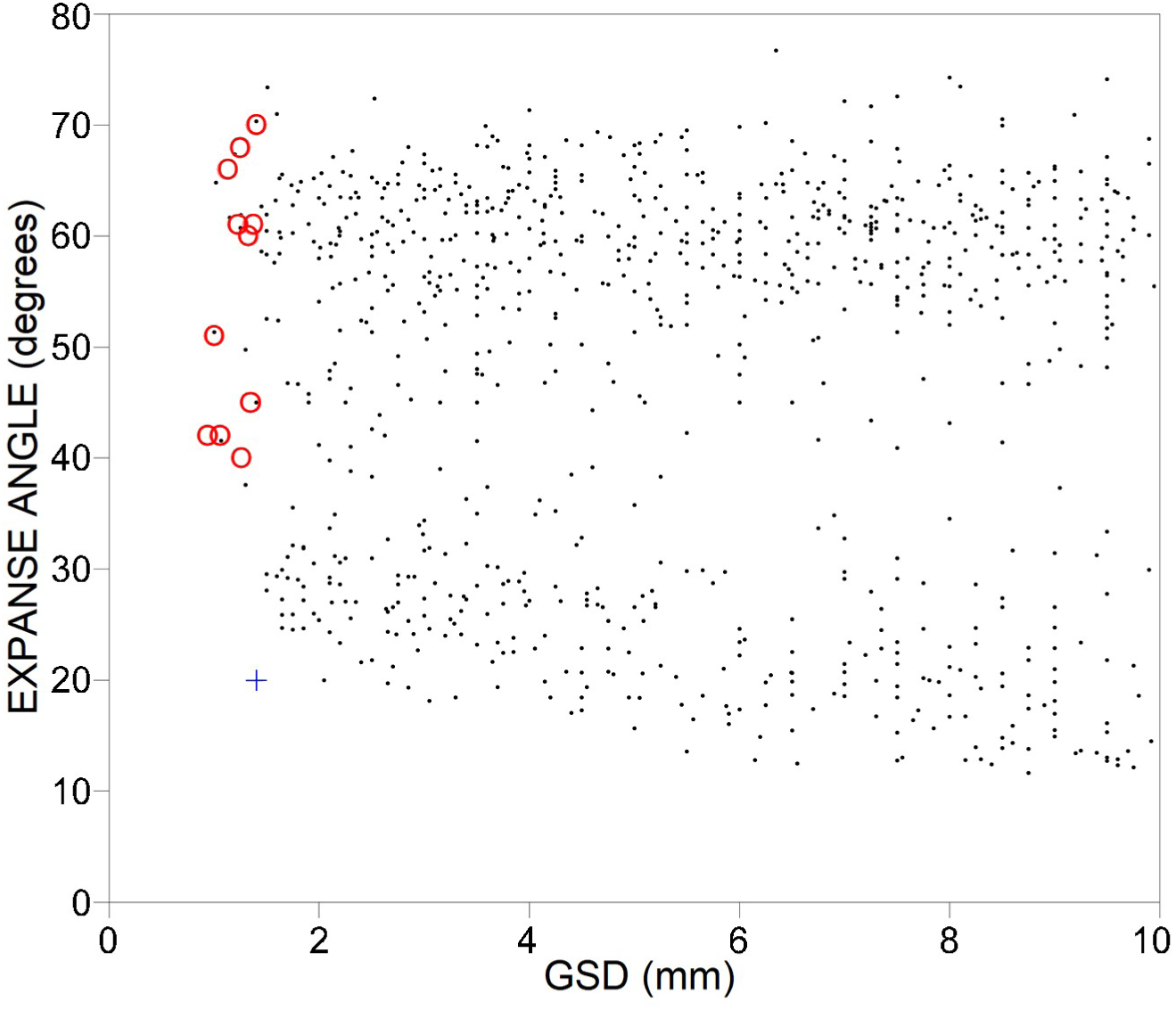
Truncated morphospace of shell shape versus size for stylommatophoran genera with mean genus GSDs < 10 mm. The superimposed points are the miniature species from Table 2 (red circles) and the type of the basommatophoran *Carychium nannodes* (+).

The concentration of the existing shell shapes in some regions of the morphospace and their absence from other regions may be explained by four types of evolutionary constraints: geometric, functional, phylogenetic and developmental (McGhee, 2007). To explain the observed increase of the LPA below the GSD of 10 mm, I will consider the first two constraints. Actually, there is no obvious geometric constraint that may prevent the formation of shells with expanse angles below the LPA. Such shells are indeed present in the Basommatophora (see below). In the next sections, I will propose and discuss two functional constraints to explain the observed inverse correlation between LPA and GSD (Fig. 3).

### Relative surface area as a possible functional constraint of shape in miniature snails

As discussed above, water loss is a potentially detrimental outcome of increased relative surface areas that accompany shell miniaturization. Consequently, evolutionary processes would be expected to reduce surface areas to decrease water loss from exposed surfaces. The relative surface area of an object depends not only on its dimensions (or volume) but also on its shape, a sphere having the smallest relative surface area for a given volume (Schmidt-Nielsen, 1984). Likewise, it can be demonstrated, by using the appropriate volume and surface area equations, that a cylinder (or a cone) whose height and diameter are limited by a common maximum value (similar to a fixed GSD in the case of a snail shell) will have its smallest relative surface area when it is equiaxial (H = D). It is reasonable to expect that the relative surface area of a snail shell has a corresponding dependence on its shape. Therefore, as shells evolve towards smaller dimensions under environmental conditions that are conducive to water evaporation, their shapes should become less tall and more equiaxial to reduce relative areas of evaporative surfaces.

### Egg size as a possible functional constraint of shape in miniature snails

Since the egg volume cannot decrease indefinitely, there must be a minimum viable egg volume that is approached or reached at very small shell dimensions. Egg dimensions of only a handful of stylommatophoran species with shell dimensions less than about 2 mm are available in the literature. In seven species of *Vertigo* (mean shell heights, 1.75-2.3 mm), mean egg diameters were 0.54-0.78 mm (Myzyk, 2011). In *Punctum pygmaeum* (Draparnaud) (shell diameter up to about 1.6 mm) mean egg dimensions were 0.51 × 0.44 mm (Baur, 1987). I have estimated the protoconch diameters of *Angustopila fabella* Pall-Gergely & Hunyadi and *A. subelevata* from their photographs in Pall-Gergely et al. (2015) as 0.35–0.39 mm. If an estimated 0.05 mm is added to account for the thickness of the egg shell (Myzyk, 2011), the diameters of the *Angustopila* eggs would become 0.4-0.44 mm. It follows that the diameter of 0.4 mm is approximately the minimum viable egg dimension of stylommatophoran land snails. Even the mean egg diameter of the much larger *Cecilioides genezarethensis* Forcart (mean shell height, 6.6 mm) was 0.4 mm (Heller et al., 1991). It is expected that the smaller a snail becomes the more difficult it will be to fit into its shell more than one developed egg at a time (Rensch, 1959). There is evidence that the clutch size indeed becomes one at shell sizes below a GSD of about 2 mm (Myzyk, 2011; Kramarenko, 2013). These considerations conform to Rensch’s (1959) generalization that “the eggs set a lower limit to body size”. This means that the size of the smallest viable hatchling determines the size of the smallest egg, which then becomes the principal determinant of the minimum possible volume of the adult snail that is going to accommodate the egg.

These arguments imply that when a stylommatophoran lineage is evolving miniature snails, the egg diameter cannot become smaller than about 0.4 mm, which may apparently be reached even above a GSD of 1.5 mm. Upon further reduction of shell size, the egg volume relative to the shell volume begins to increase. Therefore, to accommodate the increasing fractional space taken up by the egg, the fractional contributions of all or some of the other components of the total shell volume (the snail’s body, the extra shell space and the shell material) will have to decrease until they too reach their respective limits (Appendix). Alternatively, the shell volume can be enlarged by increasing the diameter of a tall shell (ε < 40°) while keeping the shell height (GSD) the same (Appendix). However, a tall shell even with a large enough volume may still not accommodate an egg unless the diameter of the body whorl (the shell width in many shell shapes) is greater than twice the egg diameter. Only then will there be room across the shell for the egg, the snail’s body, the columella and the shell walls. If the GSD continues to decrease, further flattening of shells (enlargement of ε), which increases the diameter of the body whorl, is necessary to accommodate at least one egg. Accordingly, the smaller a shell gets the flatter it becomes even though the shell volume begins to decrease once ε exceeds 45° (Appendix). This leads to a prediction: in miniature species the expanse angles of smaller adults should be larger than those of larger adults. I have tested this prediction using the data available for three *Angustopila* species. In *Angustopila subelevata* and *A. szekeresi* Pall-Gergely & Hunyadi the expanse angles do seem to increase as the adult GSD decreases, while in the slightly larger *A. fabella*, the expanse angles appear to be independent of the adult GSD (Fig. 4).

**Figure 4.**
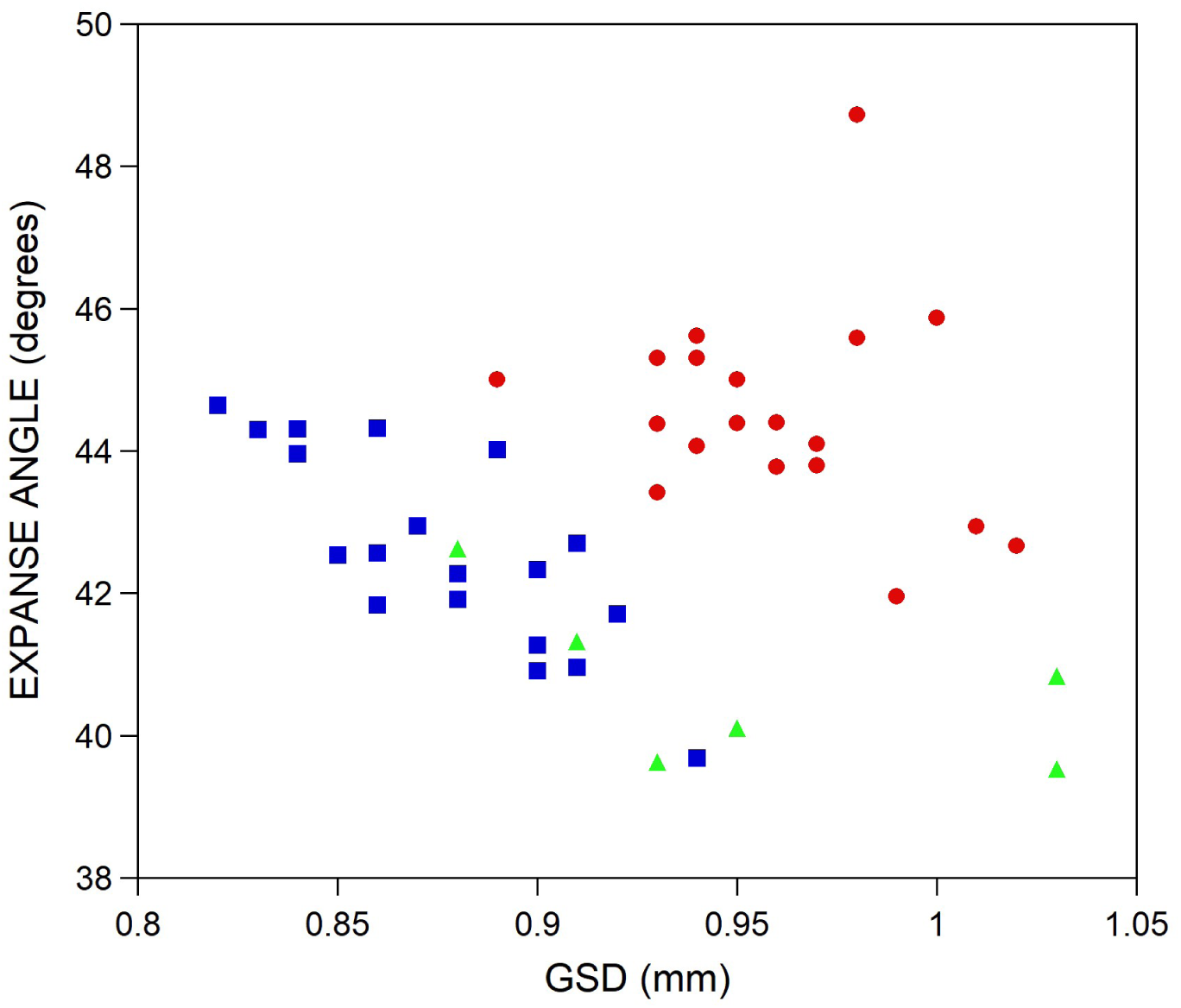
Variation of expanse angles with GSDs of the adult shells of *Angustopila subelevata* (squares), *A. szekeresi* (triangles) and *A. fabella* (circles).

An exception that strengthens this proposed constraint is given by the basommatophoran land snail *Carychium nannodes*. The expanse angle of *C. nannodes* is smaller than the LPA for the stylommatophorans having the same GSD value (Fig. 3). The relevant difference between the shells of *Carychium* and the stylommatophorans is that the adult *Carychium* resorbs the columella and the internal partitions in the spire of its shell (Pilsbry, 1948). Thus, *C. nannodes* can afford a narrow shell, because shell resorption must increase the inner volume available both for the snail and its egg (Appendix). In contrast, shell resorption has not been demonstrated in stylommatophorans (Solem, 1983).

Ontogenic changes in shell shape are often observed in the form of flat juveniles growing up into tall adults, as, for example, in the 3-mm species *Lauria cylindracea* (Heller et al., 1997). Since juveniles do not carry eggs, such shape changes may stem not only from the necessity to reduce surface areas to decrease water loss during early developmental stages, but also from phylogenetic and developmental constraints. Cain (1981) attributed ontogenic changes in shell shape–albeit in the much larger *Cerion*– to the change of the snails’ microhabitat from horizontal soil surfaces, where the flat-shelled juveniles dwell, to more vertical surfaces on which the tall-shelled adults rest.

## ACKNOWLEDGEMENTS

I thank Barna Páll-Gergely for providing me with dimensional data for *Angustopila* species and Tim Pearce for a detailed review of an earlier draft and for loaning me Schileyko (1998-2007).

## APPENDIX

The total shell volume (V_sh_) incorporates the volumes of four components: snail’s body (V_sn_) + egg (V_eg_) + extra shell space (V_es_) + shell material (V_sm_). Although the egg is within the snail’s body, here it is treated separately. The extra shell space is the empty volume between the aperture and the snail’s body when the latter has withdrawn away from the aperture for protection. The following equation expresses these volume components as fractions of V_sh_.

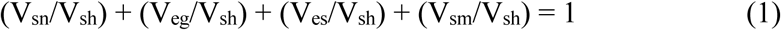

Once the egg reaches the minimum viable egg volume it cannot become smaller. Therefore, further reduction of the shell dimensions will decrease V_sh_ and increase (V_eg_/V_sh_), while necessarily decreasing the relative contributions of the other components.

The generalized volume equation for a geometric solid with two axes (height and diameter), such as a cone or a cylinder, is V = constant × H × D^2^ (Schmidt-Nielsen, 1984). This relationship is expected to hold for snail shells also, although an accurate volume cannot be calculated without knowing the value of the constant term (Örstan, 2011). It follows that as the diameter of a tall shell increases, while its GSD, which is the height at ε < 45° (Table 1), remains constant, V_sh_ increases and (V_eg_/V_sh_) decreases creating more space for the other volume components (equation 1). One caveat is that the volume of a cone or a cylinder, and probably also of a shell, reaches its maximum value when H = D (ε = 45°) and for a given GSD the shell volume will be less at all other values of ε.

Another option is to increase the space available inside a shell is by resorbing the columella and the internal partitions in the spire of the shell without necessarily thinning the external shell walls. This decreases (V_sm_/V_sh_) and increases the volume available for the other three components (equation 1). Shell resorption takes place in the adult basommatophoran *Carychium nannodes*, which has a shell narrower than that expected at its GSD value (Fig. 3).

